# Comparative analysis of *SQUAMOSA PROMOTER BINDING PROTEIN-LIKE (SPL)* gene family between bryophytes and seed plants

**DOI:** 10.1101/2023.02.27.530190

**Authors:** Alisha Alisha, Zofia Szweykowska-Kulińska, Izabela Sierocka

## Abstract

*SQUAMOSA-PROMOTER BINDING PROTEIN-LIKE* (*SPL*) genes encode plant-specific transcription factors which have been found to be conserved in green plants lineage. SPL proteins are important regulators of diverse plant developmental processes in bryophytes and vascular plants. In our study, we took advantage of available genome sequences of representatives of each bryophyte clade to investigate the relationships of *SPL* genes between bryophytes and model angiosperm *Arabidopsis thaliana*. We have identified four *SPL* genes in each of the two hornworts species, *Anthoceros agrestis* and *Anthoceros punctatus*, what is similar to the set of *SPL* genes present in the liverwort *Marchantia polymorpha*. Thus, the analyzed hornworts and liverwort genomes encode a minimal set of *SPL* genes in comparison to other land plants that may resemble an archetype of *SPL* genes present in the ancestor of land plants. The phylogenetic analysis revealed the presence of four *SPL* groups. Comparative gene structure analysis showed that *SPLs* share similar exon-intron organization within the same phylogenetic group with some exceptions in hornworts. While we have identified conserved protein motifs between bryophytes and *Arabidopsis* in three out of four phylogenetic groups, the motif content differed explicitly in the fourth group. Since current understanding of *SPL* genes mostly arises from seed plants, the presented comparative and phylogenetic analysis will provide better understanding of *SPL* gene family from the representatives of the oldest living land plants.

## Introduction

Transcription factors (TFs) are DNA-binding proteins that specifically regulate gene transcription activation or repression. *SQUAMOSA PROMOTER BINDING PROTEIN-LIKE* (*SPL*) genes encode plant-specific transcription factors that are widely distributed from unicellular algae to angiosperms (Chao *et al*., 2017). For the first time they were described in snapdragon based on their ability to specifically bind to the promoter of floral meristem identity gene *SQUAMOSA*, which encodes a MADS-box transcription factor (Klein *et al*., 1996). SPL proteins are diverse in their primary protein structures but share characteristic SBP domain (SQUAMOSA-PROMOTER BINDING PROTEIN). The SBP domain is composed of highly conserved 76 – 78 amino acid residues consisting of two zinc-binding sites, Cys-Cys-Cys-His and Cys-Cys-His-Cys, respectively. The zinc ions present in the SBP domain are crucial for its proper folding and stability which is required for recognition and binding to specific DNA sequences. (Yamasaki *et al*., 2004; Birkenbihl *et al*., 2005). Additionally, a bipartite nuclear localization signal (NLS) motif resides at the C-terminal end of the SBP domain which overlaps with the second zinc-binding site. This NLS is required for the nuclear import of SPL proteins (Birkenbihl *et al*., 2005; Zhang *et al*., 2019).

In the model plant *Arabidopsis thaliana*, 16 members in the *SPL* family were identified, whereas in moss, *Physcomitrium patens*, 13 members were found (Cardon *et al*., 1999; Riese *et al*., 2007, 2008). With the progress of sequencing techniques, the identification and evolution of the *SPL* gene family has been widely investigated in angiosperms. Genomic sequencing has revealed 19, 28, 31, and 56 *SPL* genes in *Oryza sativa, Populus trichocarpa, Zea mays* and *Triticum aestivum*, respectively, indicating dynamics of the *SPL* genes evolution within angiosperms (Hultquist and Dorweiler, 2008; Miura *et al*., 2010; Li and Lu, 2014; Zhu *et al*., 2020). However, for the other land plant lineages these studies are heavily underrepresented with only one moss, *Physcomitrium patens, SPL* gene family being used in comparative analyses (Preston and Hileman, 2013; Zhang *et al*., 2015). Based on the length of encoded proteins, the *SPL* genes can be divided into two classes. First, contains genes encoding proteins longer than 800 amino acids (ex. At*SPL1, 7, 12, 14*, and *16*) while second groups genes encoding proteins shorter than 800 amino acids (the remaining 12 At*SPL* genes) (Cardon *et al*., 1999; Birkenbihl *et al*., 2005; Preston and Hileman, 2013; Zhang *et al*., 2015). Phylogenetic studies have shown that increase in the *SPL* gene number during the evolution of land plants was mainly the result of expansion of genes with 2–10 exons encoding shorter proteins. Moreover, only within this group of *SPLs* the expression of a large number of genes is regulated by the miR156 and miR529 family members through mRNA cleavage and/or translational repression (Zhang *et al*., 2015). In *Arabidopsis*, miR156 targets ten members of the *SPL* family to regulate different biological processes including vegetative-to-reproductive phase transition, abiotic stress responses and lateral organs development (Yamaguchi *et al*., 2009; Shikata *et al*., 2009; Jung *et al*., 2012). Whereas in moss *P. patens*, only transcripts of three *SPL* genes are recognized by miR156. Studies have shown that the deletion of one of them, Pp*SBP3*, accelerates and increases the number of developing leafy buds from the juvenile protonemal phase, showing that in the wild type plant Pp*SBP3* acts as a negative regulator of moss phase-transition from tip-growing protonema to leafy gametophores (Cho *et al*., 2012). Although not directly comparable due to the life cycle differences of mosses and angiosperms, this function is somewhat similar to At*SPL14*, which functions to delay the transition to adult development (Stone *et al*., 2005).

miR156 is one of the few highly evolutionarily conserved miRNAs in plants (Pietrykowska *et al*., 2022). However, miR156 is lacking in the microtranscriptome of liverwort, *Marchantia polymorpha*. Instead, miR529 is present as an equivalent module which regulates the transcript level of one of four *SPL* genes, Mp*SPL2* (Lin et al., 2016; Tsuzuki et al., 2019). Recently it was shown that miR529c loss-of-function or miRNA-resistant version of Mp*SPL2* produces functional reproductive organs even in the absence of far-red light, the environmental signal required for the transition from vegetative to reproductive phase in the wild-type plants. Interestingly, Mp*SPL2* knock-out plants can produce gametangiophores under proper environmental conditions. This data suggests that the miR529-Mp*SPL2* module is important for the reproductive transition under far-red light conditions, but it is not essential for gametangia development. Similar to the role of miR156-*SPL* module in seed plants, the miR529c-Mp*SPL2* module was found to regulate the reproductive transition in *M. polymorpha* (Tsuzuki *et al*., 2019).

Although the *SPL* gene family has been widely studied in many species, research on the classification and evolution of *SPLs* is still missing in liverworts and hornworts which represent the ancient lineages of land plants. In our study, we took advantage of available genome sequences from representatives of each bryophyte clade to investigate the relationships of *SPL* genes between bryophytes and angiosperms. The *SPL* gene and protein sequences for *M. polymorpha, P. patens* and *A. thaliana* were downloaded from each of the plant genome database according to their annotation (‘Phytozome v13’, ‘TAIR - Home Page’; Bowman *et al*., 2017). For hornworts, *Anthoceros agrestis* and *Anthoceros punctatus*, available genomic resources were used to identify and annotate the *SPL* gene family members (Szövényi *et al*., 2015; Li *et al*., 2020). Subsequently, we performed phylogenetic analysis to trace the evolutionary relationships between SPL proteins from bryophytes and *Arabidopsis*. Furthermore, we comprehensively analyzed features of the conserved SBP domain and additional conserved protein motifs distribution in each studied clade, followed by exon-intron gene structure analysis. Moreover, the availability of expression data from RNA-sequencing experiments for *A. thaliana, P. patens, M. polymorpha* and *A. agrestis* allowed us to investigate the expression profiles of *SPL*s in these species. Our study provides substantial insights into the understanding of the origin and evolution of the *SPL* gene family in embryophytes and emphasizes the importance of studying the biological relevance of *SPLs* in the representatives of bryophytes.

## Materials and Methods

### Identification of *Anthoceros SPL* genes and bioinformatic analysis

Genomes with available annotation of two hornwort species, *Anthoceros agrestis* (Bonn) and *Anthoceros punctatus* were downloaded from University of Zurich database (‘UZH -Hornworts’). The protein sequences of *A. thaliana, P. patens* and *M. polymorpha* were retrieved from the *Arabidopsis* information resource database TAIR version 10, (‘TAIR - Home Page’), Phytozome version 13 (‘Phytozome v13’; Goodstein *et al*., 2012) and MarpolBase database, respectively (Bowman *et al*., 2017). These included 16 *Arabidopsis*, 13 *Physcomitrium* and four *Marchantia* SPL protein sequences which were used as queries to identify putative SPL protein sequences from *A. agrestis* and *A. punctatus* by using local BLASTp (Table S1). An e-value of <10^−5^ and bit-score >100 was used as an initial cut-off to claim significant matches, remove redundant hits and select unique sequences for further analysis. In order to ensure the presence of SBP domain, all the candidate SPLs were searched against SMART (Letunic and Bork, 2018) and ScanProsite databases (de Castro *et al*., 2006).

The miRNA binding sites were identified in *Anthoceros SPL* gene transcripts using psRNATarget server (Dai and Zhao, 2011). The molecular weight (Mw) and theoretical isoelectric point (p*I*) of *Anthoceros* SPL protein sequences were calculated using Compute pI/Mw tool in the ExPASy server (‘SIB Swiss Institute of Bioinformatics’; Gasteiger *et al*., 2005). The subcellular localization was predicted online by WoLFPSORT (‘Website’; Horton *et al*., 2007).

### Phylogenetic tree construction

In order to identify phylogenetic relationships between SPL proteins from each bryophyte clade representative and dicots representative, full length protein sequences from *A. agrestis, A. punctatus, M. polymorpha, P. patens* and *A. thaliana* were aligned using CLUSTALW tool in MEGA11 (Tamura *et al*., 2021). Further, the phylogenetic tree was constructed by using bootstrap maximum likelihood method with 1000 replicates to obtain support values for each branch. CRR1 protein from algae *Chlamydomonas reinhardtii* was used as an outgroup (Kropat *et al*., 2005; Sommer *et al*., 2011).

### Gene structure analysis and conserved protein motifs characterization

The exon-intron structures of *SPL* genes were analyzed by GSDS software (Hu *et al*., 2015). Conserved motif analysis in SPL proteins was performed using MEME program (Multiple EM for Motif Elicitation’ v5.4.1) (Bailey *et al*., 2015). The number of predicted motifs was set to 20 with the default parameters (minimum width 6 and maximum width 50). The SBP domain sequences were aligned using CLUSTALW in Jalview software (Waterhouse *et al*., 2009).

The protein sequences annotated as SPL proteins from *C. reinhardtii* were downloaded from Phytozome version 13 (‘Phytozome v13’) (Table S1). Further, all putative *Chlamydomonas* SPL sequences were queried against SMART (Letunic and Bork, 2018) and ScanProsite databases (de Castro *et al*., 2006) to confirm the conserved SBP domain presence. Only ten sequences containing the conserved two zinc-binding sites, Cys-Cys-Cys-His and Cys-Cys-His-Cys, were selected for further analysis. The sequence logo for SBP domain sequences was generated by WebLogo 3 platform (Crooks *et al*., 2004).

### Cis-acting element analysis of *SPL* gene transcripts

The 1500 bp upstream sequences from the start codon for each *SPL* gene sequences from *M. polymorpha, P. patens* and *A. thaliana* were retrieved from the respective genomic resources. For the two *Anthoceros* species bedtools were used to retrieve 1500 bp upstream sequences for each *SPL* gene (Quinlan and Hall, 2010). The putative *cis*-elements were identified using PlantCARE software (Lescot, 2002). The identified motifs shown to be putatively involved in plant growth and development, light responsiveness, stress and phytohormone responses are summarized in this study (Table S4).

### Expression profiling of *SPL*s from different developmental stages in *Arabidopsis* and Bryophytes

The expression data for *A. thaliana* and *P. patens* were downloaded from expression atlas, EMBL-EBI and PEATmoss database, respectively (Liu *et al*., 2012; Ortiz-Ramírez *et al*., 2016; Fernandez-Pozo *et al*., 2020). The expression data for *M. polymorpha* and *A. agrestis* were downloaded from (Kawamura *et al*., 2022) and (Li *et al*., 2020), respectively. The detailed description of the RNA-seq datasets used in our analysis is provided in Table S5. A heat map presenting the expression profiles of *SPL* genes for each plant was generated using RStudio (‘Posit’, 2022).

## Results

### Identification of *SPL* genes from hornworts

BLASTP was used to identify the *SPL* genes in two hornwort genomes, *A. agrestis* and *A. punctatus*, while SMART and ScanProsite tools were used to validate the results (Sigrist *et al*., 2013; Letunic *et al*., 2021). After removing the redundant sequences and sequences with incomplete SBP-box domain, four *SPL* genes were identified in each of the two *Anthoceros* genomes. Nomenclature of the identified genes was carried out on the basis of their identity with the respective four members of *Marchantia SPL* family (Tsuzuki *et al*., 2016). The final *SPL*s from *A. agrestis* and *A. punctatus* were named Aa*SPL1-4* and Ap*SPL1-4*, respectively (Table 1).

**Table 1.**
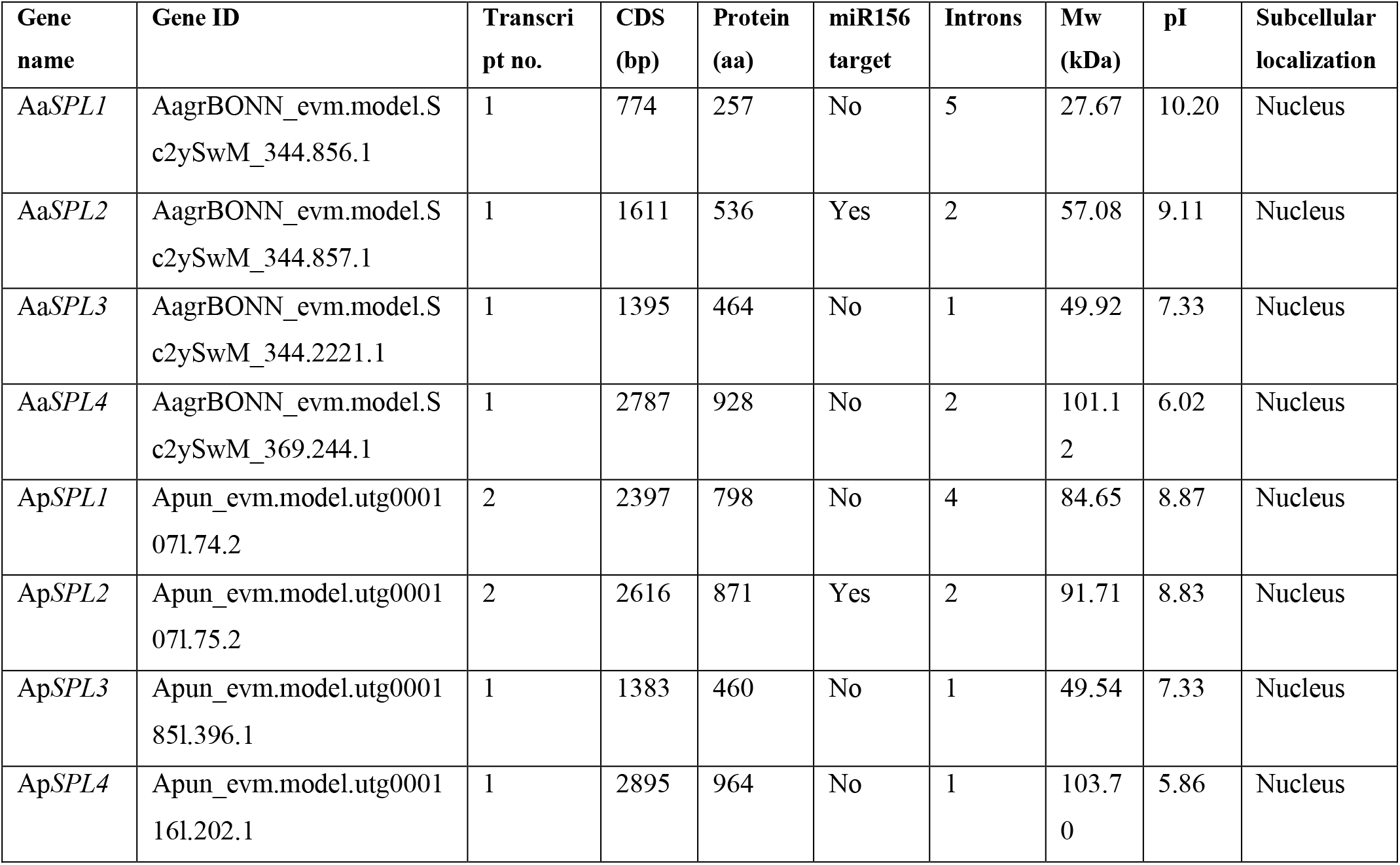
Sequence and molecular features of *SPL* genes identified in two *Anthoceros* species. Aa – *Anthoceros agrestis*, Ap – *Anthoceros punctatus*.

The number of introns varied between identified genes and ranged from 1 to 5 (Table 1). The lengths of CDS sequences varied from 774 to 2895 bp while their protein lengths varied from 257 to 964 amino acids. The molecular weight of deduced SPL proteins ranged from 27.7 to 103.7 kDa while their isoelectric points ranged from 5.86 to 10.20. The subcellular localization of both *Anthoceros* SPL proteins was predicted to be in the nucleus. These results have shown the diversity within structural features of *SPL* genes as well as in physical and chemical properties of SPL proteins in two *Anthoceros* species.

### Comparative evolutionary analysis of *SPL* gene family between bryophytes and land plants

To evaluate the evolutionary relationships of SPL proteins from seed plants and bryophytes, phylogenetic analysis was conducted using full length SPL protein sequences from dicot *A. thaliana*, moss *P. patens*, liverwort *M. polymorpha* and two hornworts *A. agrestis* and *A. punctatus*. Based on the obtained phylogenetic tree, SPL proteins were classified into four distinct groups, Group 1 – Group 4 (Fig. 1), where at least one member from the analyzed plant species was assigned to each group. *M. polymorpha* and both *Anthoceros* species have only four *SPL* members as compared to 13 *SPL* genes from *P. patens* and 16 from *A. thaliana*. Up to date these are the smallest *SPL* gene families identified in land plants what may reflect the starting point of evolutionary expansion of the *SPL* gene family in land plants. Moreover, the *Anthoceros* and *Marchantia* SPL proteins from Groups 1, 2 and 4 are close to each other on neighboring branches which may indicate their common origin. The separate grouping of two pairs of paralogous proteins from *Arabidopsis*, AtSPL3/6 and AtSPL4/5 indicates that these proteins emerged after the divergence of bryophytes and angiosperms and thus, they do not have related SPL proteins in bryophyte representatives. Therefore, they were not included in any of the groups.

**Fig. 1:**
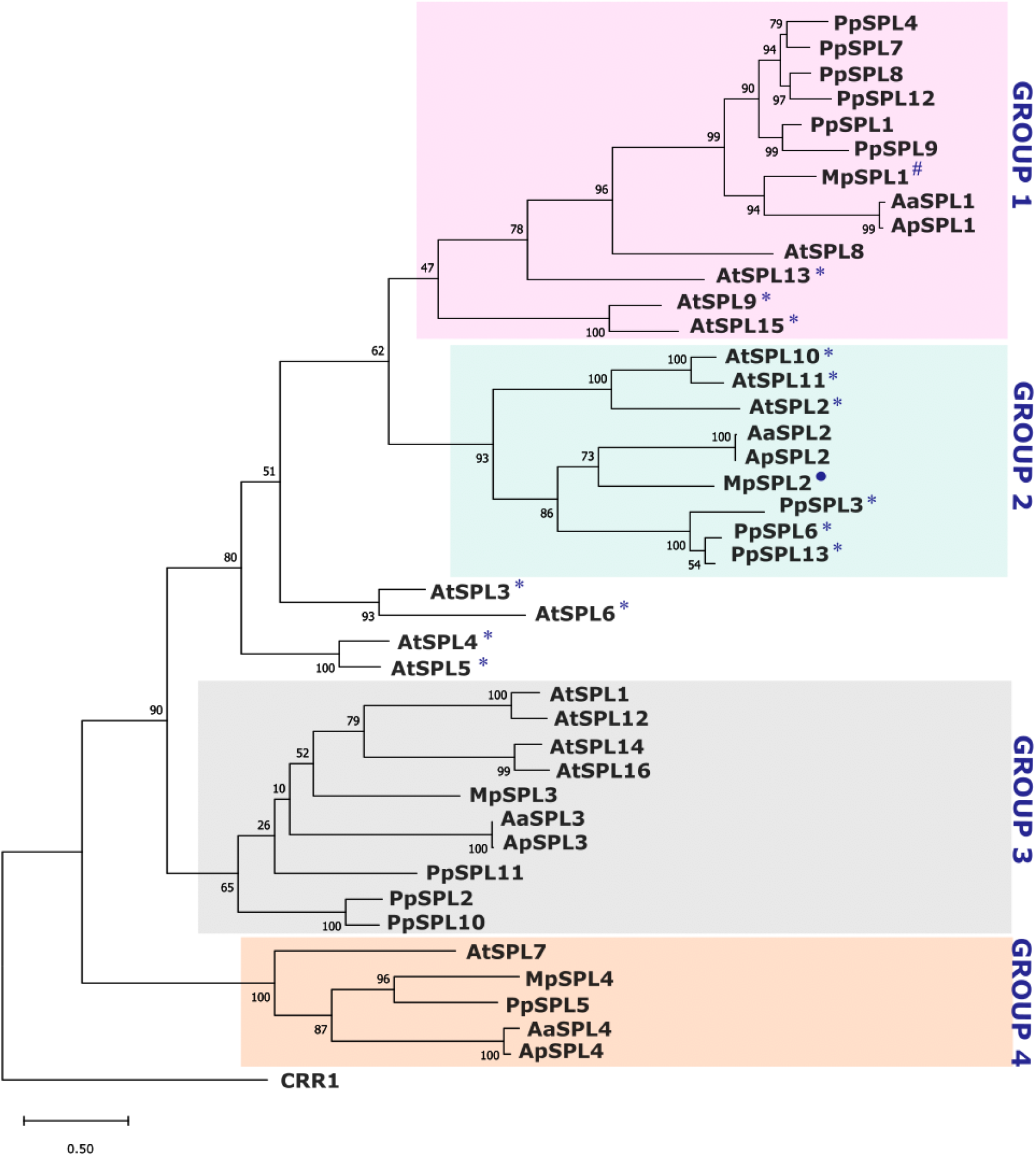
Phylogenetic relationships of SPL proteins from *A. thaliana, P. patens, M. polymorpha, A*.*agrestis* and *A. punctatus*. The tree was constructed using the maximum-likelihood method in MEGA 11 software (Tamura *et al*., 2021). Number on branches indicates the bootstrap values (%) for 1000 replications. SPL members from the same species are preceded by the prefixes: At - *A. thaliana*, Pp – *P. patens* Mp - *M. polymorpha* Aa – *A. agrestis*, Ap – *A. punctatus*. The CRR1 protein from *C. reinhardtii* was used as an outgroup to root the tree. *SPL* genes marked by *, ° and ^#^ are regulated by miR156, miR529c and Mpo-miR13, respectively.

For many plant species, it was shown that among the *SPL* family, some of the members undergo post-transcriptional gene expression regulation by conserved miRNAs, miR156 or miR529 and *Marchantia* specific Mpo-miR13 (Streubel *et al*., 2023). As no experimental data are available for *A. agrestis* and *A. punctatus* microtransriptomes, we applied homology-based search to identify miRNA candidates which could target *Anthoceros SPL* gene transcripts. Mature miRNA sequences from liverwort *Pellia endiviifolia* (miR156) and *M. polymorpha* (miR529c and Mpo-miR13) were used as an input sequences (Alaba *et al*., 2015; Lin *et al*., 2016; Tsuzuki *et al*., 2019; Streubel *et al*., 2023). Since we did not find any sequences matching to miR156/529c or Mpo-miR13 in both *Anthoceros* genomes, we used *Anthoceros SPL* transcript sequences to predict potential target sites which could be recognized by these miRNAs by using psRNATarget server. Applying a stringent cut-off threshold (maximum expectation from 0 to 2) which reduces the false positive predictions, only *AaSPL2* and *ApSPL2* mRNAs were recognized as potential targets for miR156 and miR529c (Table S2). However, further experiments are needed to investigate the presence of miRNAs in *A. agrestis* and *A. punctatus* that could regulate Aa*SPL2* and Ap*SPL2* transcripts level. Since *SPL* genes belonging to Group 1 are also targeted by miRNAs, miR156 in *A. thaliana* and liverwort-specific Mpo-miR13 in *M. polymorpha*, we cannot rule out the possibility that in the two *Anthoceros* genomes a hornwort-specific miRNA is present to regulate the *AaSPL1* and *ApSPL1* gene transcript levels. However, future detailed microtranscriptome studies are needed to verify this hypothesis.

Group 4 was found to be somewhat distinctive from the other three groups because it was represented by single gene members from all species included in this study. Interestingly, the characteristic feature of the encoded proteins belonging to this group is the presence of a different signature C4 motif at the first zinc finger structure (Zn-1) in the SBP domain as compared to the canonical C3H motif found in all other SPL proteins. Stable number of genes in the Group 4 indicates their highly conserved character and resistance to expansion during SPL family evolution.

### Gene structure analysis of SBP-box genes

To learn about the structural diversity of *SPL* genes between bryophytes and *Arabidopsis*, we performed exon-intron structure analysis. Variations in the number and lengths of exons and introns were observed in each *SPL* clade (Fig. 2). The highest diversity in the gene exon-intron structure was observed in Group 1, as *Marchantia* and *Arabidopsis* genes contain two introns, *Anthoceros* four to five introns and *Physcomitrium* six to seven introns. Also, within this group of *SPL* genes, the position of CDS sequence encoding SBP domain is highly conserved and interrupted by an intron. On the other hand, the genes present in Group 2 showed the highest similarity between their gene structures with all other genes containing two to three introns. Additionally, all members belonging to this group also exhibit a highly conserved position of the CDS sequence encoding SBP domain i.e, between exons 1 and 2, interrupted by an intron.

**Fig. 2:**
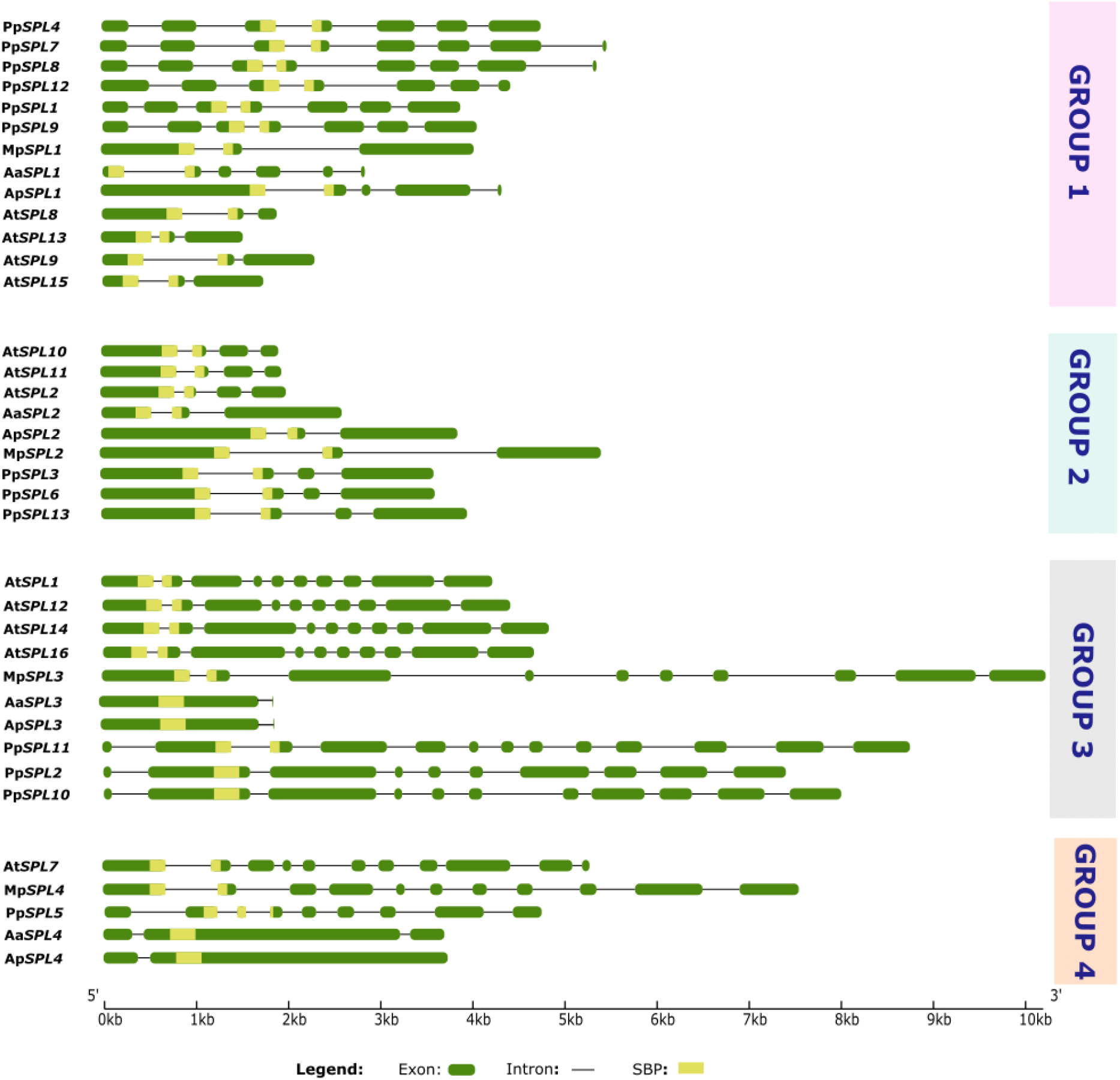
Diagram of exon-intron organization of the *SPL* gene family from *A. thaliana, P. patens, M. polymorpha, A. agrestis* and *A. punctatus*. The gene structures were analyzed using gene structure display server 2.0 (Hu *et al*., 2015) and grouped based on their phylogenetic relationships. In each gene model, exons are shown as green boxes, introns as black lines and SBP-box as yellow rectangular shading. The scale shown at the bottom represents gene lengths in base pairs.

The members belonging to Group 3 and Group 4, with the exception of *Anthoceros SPLs*, showed the highest number of introns, from eight to ten. The hornworts genes however possess only one or two introns in these phylogenetic groups. In most of the members from Group 3 and Group 4, their CDS sequence for SBP domain is interrupted by an intron with the exception of Pp*SPL2*, Pp*SPL10* and *Anthoceros SPL*s. On the other hand, the genes present in Group 2 showed the highest similarity between their gene structures with all other genes containing two to three introns. Additionally, all members belonging to this group also exhibit a highly conserved position of the CDS sequence encoding SBP domain interrupted by an intron.The highest diversity in the gene exon-intron structures was observed in Group 1, as *Marchantia* and *Arabidopsis* genes contain two introns, *Anthoceros* four to five introns and *Physcomitrium* six to seven introns. Also, within this group of *SPL* genes, the position of CDS sequence encoding SBP domain is highly conserved and interrupted by an intron.

Based on the identified exon-intron structures of *SPL* genes, differences in the intron lengths were observed between the studied species. To validate these differences, we calculated the average intron lengths of the *SPL* genes for each species. The obtained values for *Arabidopsis, Physcomitrium, Marchantia* and *Anthoceros SPL* genes were 51 bp, 156 bp, 275 bp and 104 bp, respectively showing that *Arabidopsis* and hornworts *SPL* genes possess the shortest introns, while *Marchantia* exhibits the longest introns from all the analyzed *SPL* genes. These data coincide with the data published for the genome of each plant studied, where the average intron lengths were calculated to be 164 bp in *Arabidopsis*, 278 bp in moss *Physcomitrium*, 392 bp in liverwort *Marchantia* and 104/103 in hornworts (Swarbreck *et al*., 2008; Lang *et al*., 2008; Bowman *et al*., 2017; Li *et al*., 2020).

### Identification of conserved motifs in SPL proteins

To analyze the diversity and similarity between SPL protein structures from bryophytes and *Arabidopsis*, conserved domains and motifs were identified using MEME online tool (Bailey *et al*., 2015). The co-ordinates and sequences of SBP-box domains within each SPL protein were obtained using Pfam 35.0 database (Mistry *et al*., 2021). A conserved SBP domain was found in all SPL members, represented by Motifs 2, 1 and 4 after MEME analysis (Fig. 3). Additionally, several conserved motifs were also present in the proteins belonging to the same phylogenetic group (Fig. 3). For example, Motifs 16 – 20 are group-unique motifs found only in members of *Physcomitrium* Group 1 proteins, indicating that these motifs might be important for controlling some moss lineage specific processes. Based on the protein length, Group 1 can be further divided in two subgroups: i) longer proteins represented by all *Physcomitrium* Group 1 SPL proteins, *Marchantia* MpSPL1 and *A. punctatus* ApSPL1 and ii) shorter proteins with *Arabidopsis* Group 1 SPL proteins and *A. agrestis* AaSPL1. Although similar in length to moss proteins, ApSPL1 and MpSPL1 do not exhibit characteristic arrangement of additional motifs identified in *Physcomitrium* Group 1 members. ApSPL1 and MpSPL1 share only one motif, Motif 9, localized in the N-terminal part, upstream to the SBP domain. The hornwort ApSPL1 protein possesses one additional motif, Motif 16, downstream to the SBP domain, which is also present in moss Group 1 SPL proteins. The origin from the same phylogenetic branch and the presence of similar motifs between ApSPL1 and MpSPL1, might indicate the similarity of their biological functions. However, functional studies are needed in the Anthoceros *SPL* gene family to come to this conclusion.

**Fig. 3:**
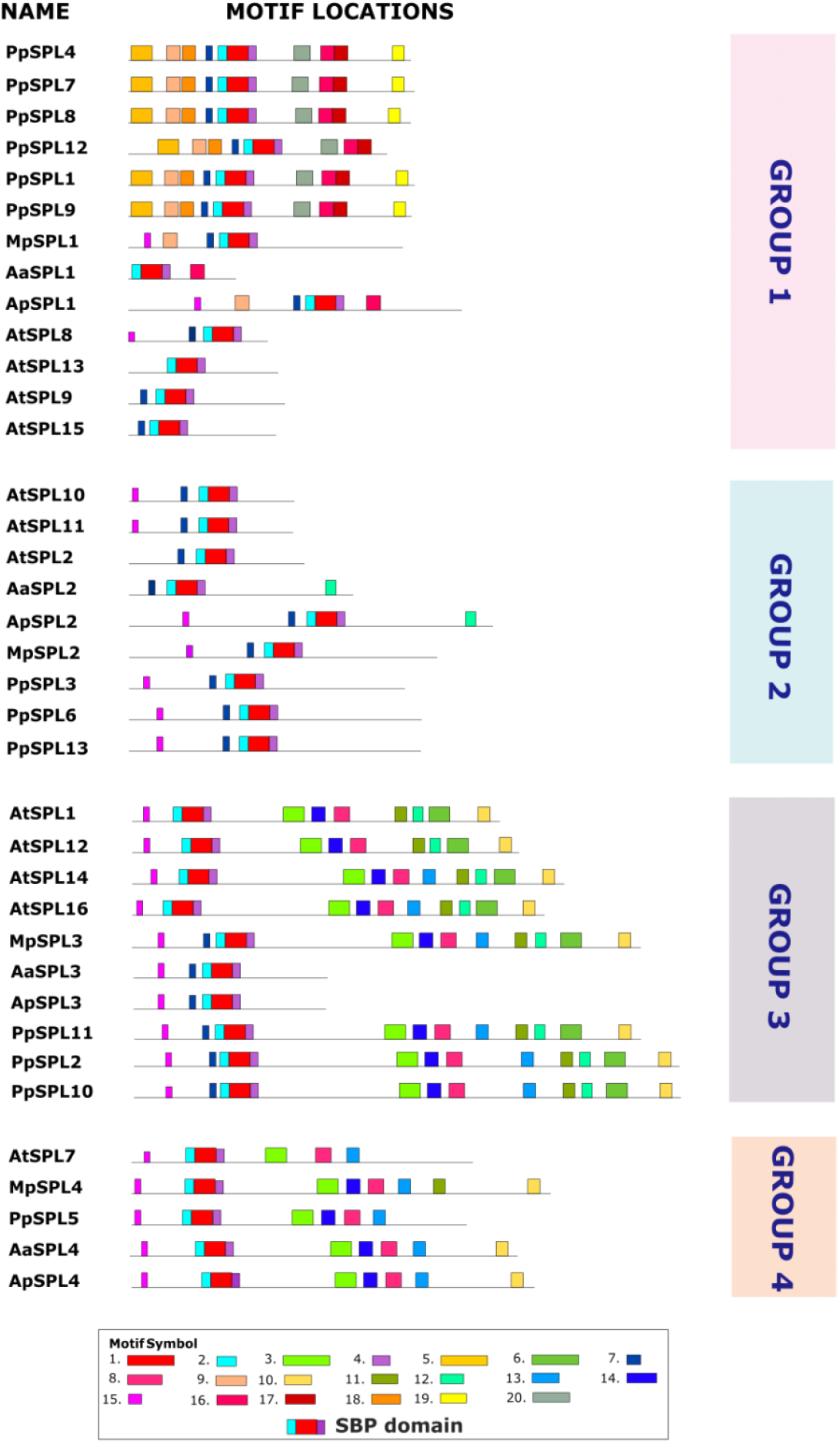
Conserved motifs in SPL proteins from *A. thaliana, P. patens, M. polymorpha, A. agrestis* and *A. punctatus*. The motif search was performed using MEME online tool (Bailey *et al*., 2015) with full length protein sequences as a query. SPL proteins are grouped according to their phylogenetic relationships. Different motifs are represented with colors shown in the legend. Motifs 1, 2 and 4 with red, blue and violet color denote SBP-box domain which is conserved amongst all SPL proteins. The consensus sequence of each motif is presented in Table S3.

The highest number of motifs were found amongst Group 3 members, while there was a relatively lower number of motifs in Group 2. With an exception of *Anthoceros* SPLs, all members belonging to Group 3 contained seven conserved motifs (Motifs 3, 14, 8, 11, 12, 6, 5). Interestingly, Motif 6 is composed of ankyrin repeats. The ANK domain has been shown to be associated with protein-protein interactions (Michaely and Bennett, 1992). What is more, three motifs present in the Group 3 SPLs, namely Motifs 3, 14 and 8, are also present in SPL proteins from Group 4 with only Motif 14 missing in AtSPL7. The high number of similar motifs shared between SPL proteins from different plant species may indicate that these proteins can play similar roles in different plant species or they may possess similar biochemical properties.

As only the SBP domain was found to be conserved and shared between all SPL proteins from bryophytes and *Arabidopsis*, we further analyzed the evolutionary conservation of amino acid composition for SBP-box domain by using ClustalW (Fig. S1). The conservation of each amino acid residue for each plant species was visualized using Weblogo tool. In addition, a weblogo for consensus sequence of SBP domain from green algae *C. reinhardtii* was generated using its ten SPL members (Fig. 4E, Table S1). The sequence logos for the SBP domain from each plant species (Fig. 4) revealed that conserved zinc-binding amino acid residues in the two zinc finger-like structures, Zn-1 and Zn-2, and the bipartite nuclear localization signal (NLS) are highly conserved in both, green algae and land plant SPL proteins. In the case of *Chlamydomonas* SBP sequences, the amino acids from the whole region of the Zn-2 site show similar conservation when compared to land plants (Fig. 4E). However, the amino acids in the Zn-1 region are significantly less conserved with characteristic positions that differ from those observed in land plants. The green alga first zinc finger region lacks the well conserved basic amino acid residues present in land plants at positions 17 – 21 from which only arginine (at position 19) is present in *Chlamydomonas*.

**Fig. 4:**
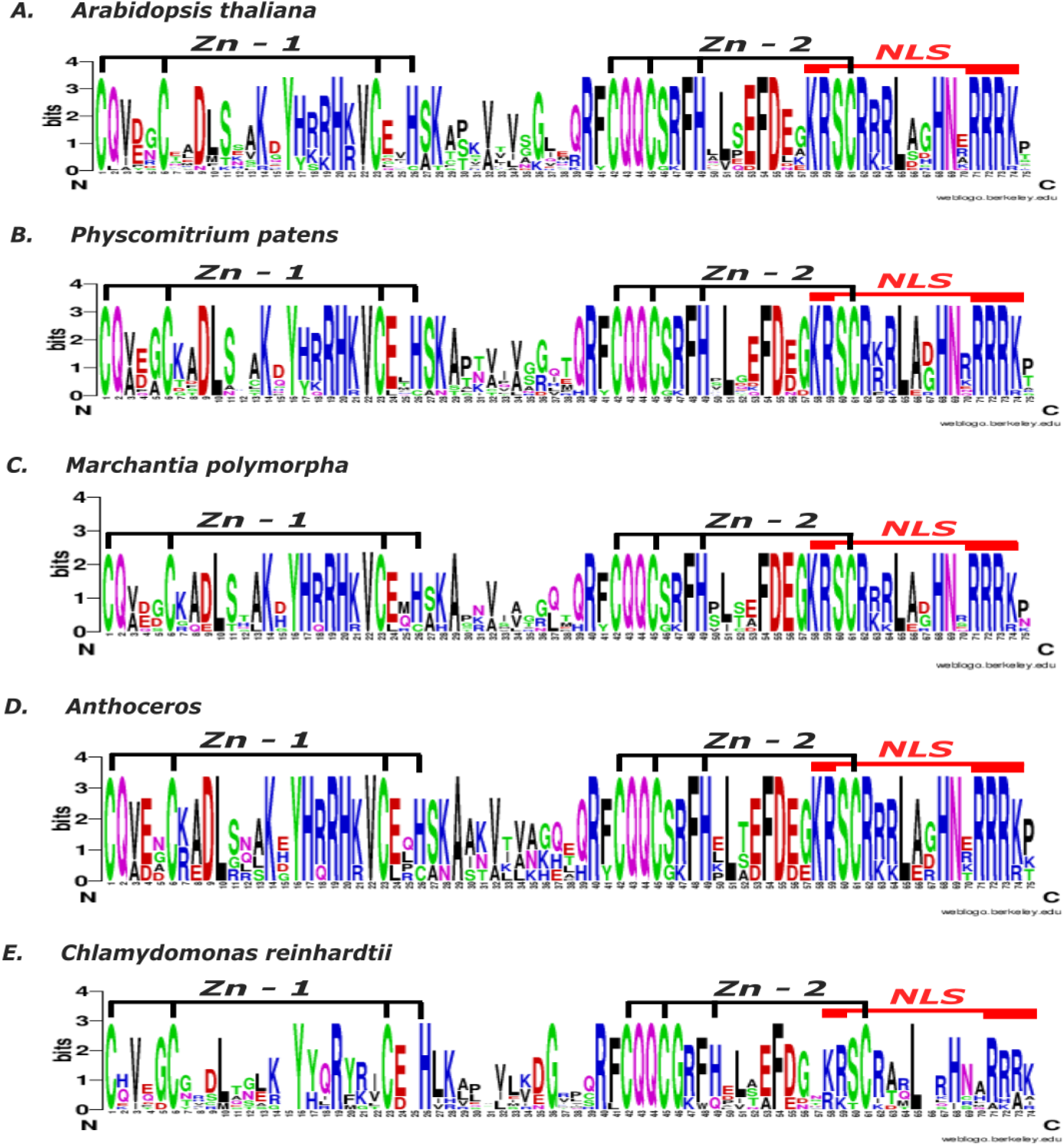
Sequence logo of conserved SBP domain of SPL proteins from (a) *A. thaliana*, (b) *P*.*patens*, (c) *M. polymorpha*, (d) *Anthoceros* and (e) *C. reinhardtii*. The sequence logo was generated using Weblogo online software (Crooks *et al*., 2004). The overall height of the stack reflects the extent of sequence conservation at that position, and the height of the letters within each stack indicates the relative frequency of each amino acid at that position.

Similarly, some divergence was also observed in the nuclear localization signal (positions 71 – 74) at the C-terminal end of the SBP domain. In general, the comparative motif analysis between land plants and green algae revealed that the SBP domain from bryophytes SPL proteins resembles that of *Arabidopsis*. Moreover, this analysis showed that the conservation of amino acids at the functional sites of the SBP domain increased during land plant SPL protein evolution. These results suggest that *SPL* genes predate the origin of land plants and the SBP domain from green algae and land plants originated from a common ancestor.

### Analysis of *cis*-elements in promoter regions of *SPL* genes

*Cis*-elements in the promoter region play important roles in the gene transcription regulation and as an adaptive mechanism to respond to different environmental conditions (Walther *et al*., 2007). To study the potential transcription regulation signals, *cis-*regulatory elements were identified in the promoter regions of investigated *SPL* genes using PlantCARE database (Table S4). A large number of *cis-*elements were detected and further classified into four subdivisions: growth and development, phytohormone response, light responsiveness and stress response (Fig. 5A and 5B, Table S4).

**Fig. 5:**
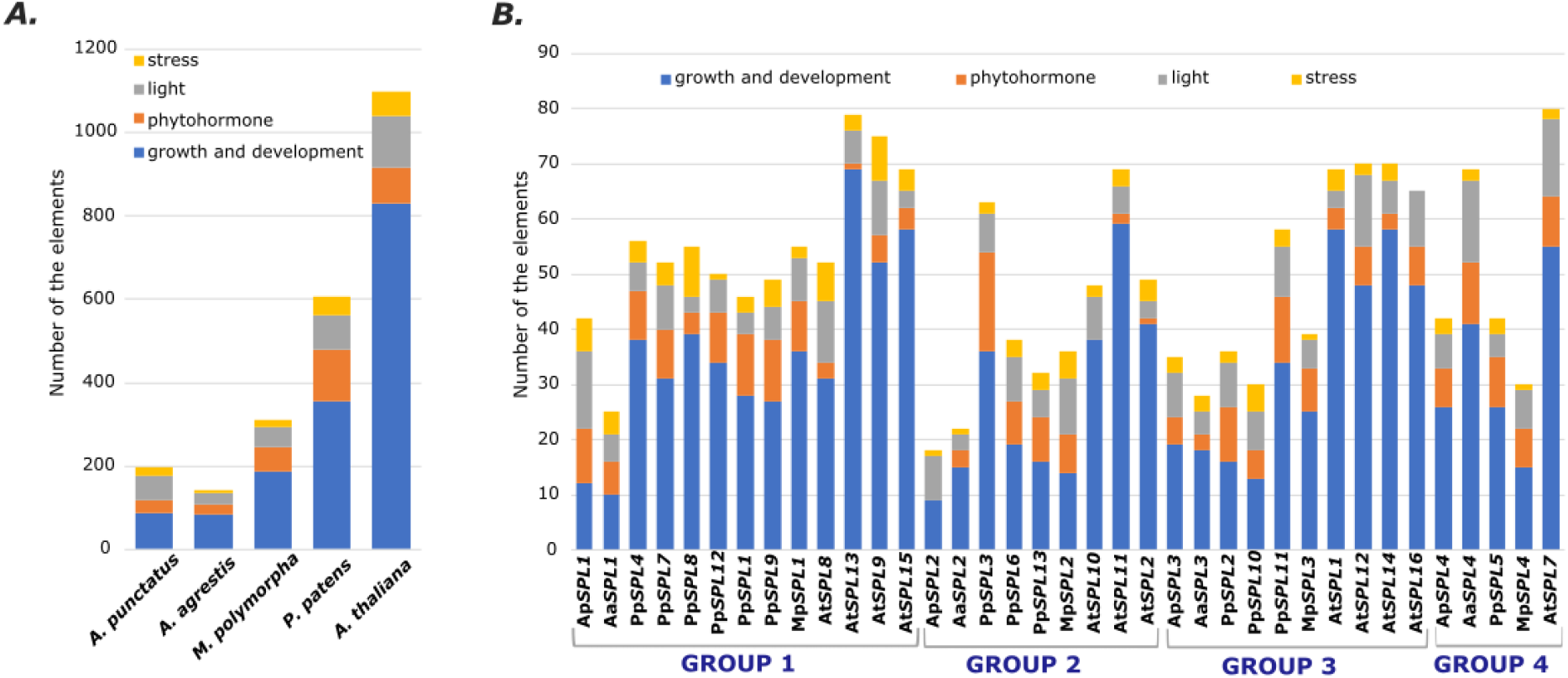
*Cis*-elements analysis of the investigated *SPL* genes from representatives of bryophytes and dicots. (A) The number of *cis*-elements in the promoter regions of *A. thaliana, P. patens, M. polymorpha, A. agrestis* and *A*.*punctatus SPL* genes. (B) The number of *cis*-elements in each *A. thaliana, P. patens, M. polymorpha, A. agrestis* and *A*.*punctatus SPL* gene promoter region grouped according to their phylogenetic relationships. The regulatory elements were detected in the 1500 bp sequences upstream of the start codon of each *SPL* gene using PlantCARE database (Lescot, 2002). The elements associated with specific functions are denoted by different colors for each gene. The detailed information concerning the *cis*-elements analysis is given in Table S4.

More than half of predicted *cis*-elements, including A-box, CAAT-box, CAT-box, CCAAT-box, GCN4 motif, NON-box, O2 site, RY element, TATA-box, AT-rich elements and circadian clock-related elements were classified under growth and development category in all studied plant species. The number of growth and development elements increased with increase in diversity of plant species. Several phytohormone responsive elements, including ABRE, AuxRR-core, CGTCA-motif, GARE-motif, TGA element, P-box, HD-Zip 3, TATC-box, TCA-element, TGA-box and TGACG-motif were identified in all four lineages. The highest number of phytohormone response elements were identified in moss and the lowest in hornworts. In the light responsive category, many elements were identified with mainly Sp1, G-box, TCT-motif and TCCC-motif being enriched. The highest number of light responsive elements were identified in *A. thaliana*. Furthermore, the identified stress response elements included ARE, TC-rich repeats, GC-motif, LTR and MBS were most common and highest in moss and dicot. In two examples, Ap*SPL2* from *A. punctatus* and At*SPL10* from *A. thaliana*, the phytohormone responsive elements were not detected (Fig. 5B). Also, the absence of stress response elements in the promoter region of At*SPL16* was observed. These results showed that *SPL* genes from different phylogenetic groups and plant species possibly participate in diverse physiological processes, developmental regulation, and abiotic stress responses.

### SPL expression profiling across different tissues in *Arabidopsis* and Bryophytes

To have a general overview about the tissue-specific expression profile of *SPL* genes in *Arabidopsis* and bryophytes representatives, we gathered the publicly available RNA-seq data for the investigated plant species from different developmental stages and organs to dissect the information about the transcript levels for each *SPL* gene (Table S5). For hornworts expression profile, only RNA-seq data for *A. agrestis* was available and used in our analysis (Li *et al*., 2020). The detected expression levels were plotted as heat maps for each plant species (Fig. 6).

**Fig. 6:**
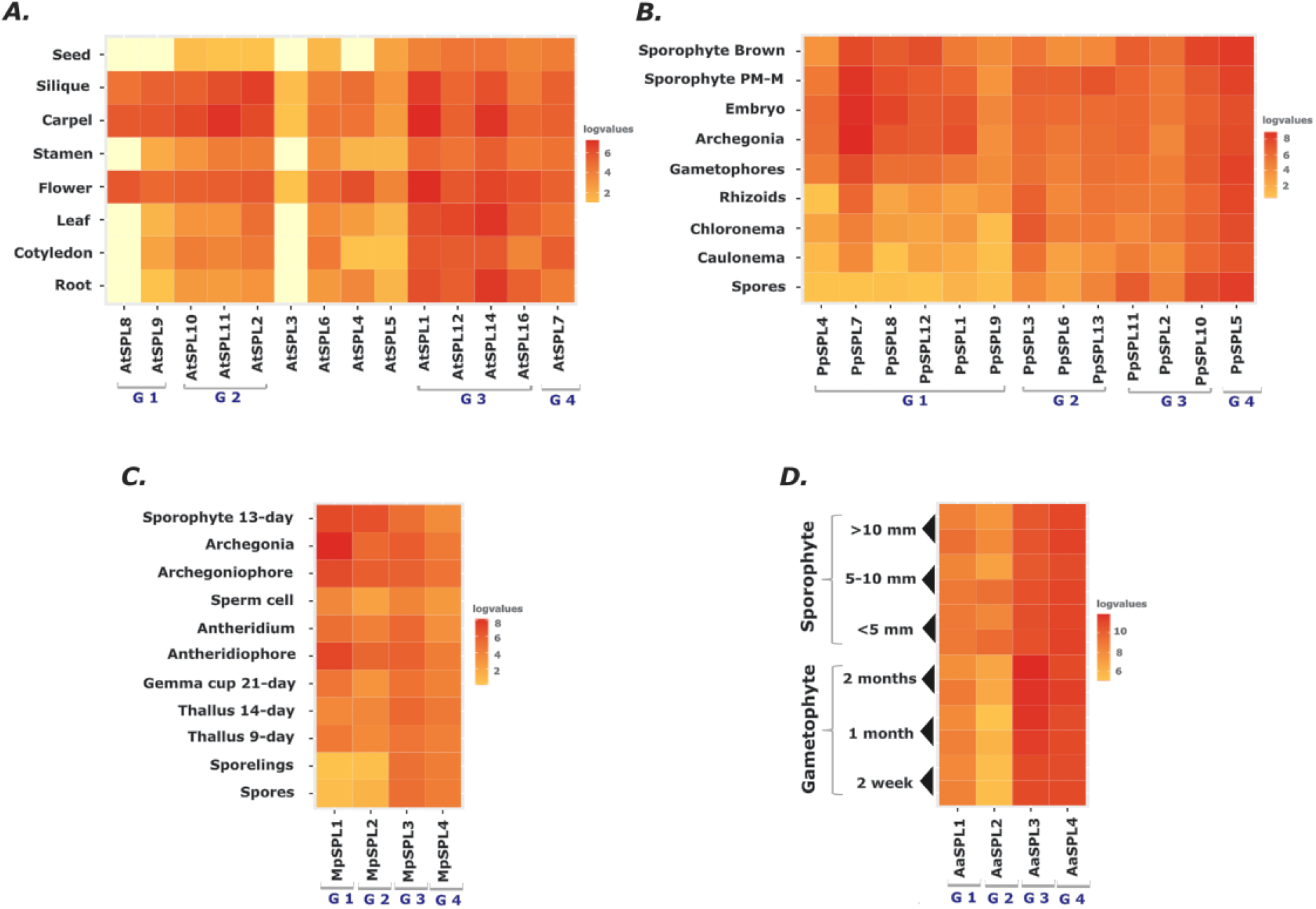
The expression profiles of *SPL* genes from different developmental stages and organs of *A. thaliana, P. patens, M. polymorpha* and *A. agrestis*. TPM and FPKM values were identified from RNA-seq data and normalized by log2 transformation for: (A) *A. thaliana* (Moreno et al., 2022), (B) *P. patens* (Fernandez-Pozo *et al*., 2020), (C) *M. polymorpha* (Kawamura et al., 2022) and (D) *A. agrestis* (Li et al., 2020). The heatmap was generated in RStudio (‘Posit’, 2022). G1-G4 denotes the names of *SPL* phylogenetic groups, Group 1-4. The red, blue and white color denotes high, low and no expression values. The white colour in the heatmap denotes no expression values in the dataset.

In the case of *A. thaliana*, 14 out of 16 *SPL* genes were expressed in the selected developmental stages (Fig. 6A). Two members belonging to Group 1, At*SPL13* and At*SPL15*, were not detected. According to experimental data showing the expression of AtSPL13 and AtSPL15 fusion proteins tagged with β-glucuronidase in transgenic plants, both these proteins accumulate at very low levels for a short time during leaf development and early stages of inflorescence development, respectively (Xu *et al*., 2016). Most probably such specific expression profiles observed for AtSPL13 and AtSPL15 proteins might be the cause that both these genes are missing in the presented analysis. In *Arabidopsis*, the expression patterns of different genes in the same phylogenetic group were observed to be rather similar, suggesting the involvement of *SPL* paralogs in the regulation of similar processes. The most specific expression pattern was observed for Group 1 and correlated mostly with flower development. The *Arabidopsis* Group 2 *SPL* genes, although expressed in more developmental stages in comparison to Group 1 *SPL* genes, also exhibited in general enriched expression during flower organs development (Fig. 6A). In turn, the At*SPLs* expression levels from Group 3 and Group 4 were high and at rather similar levels in the analyzed organs and developmental stages.

In general, based on their expression pattern, *Arabidopsis SPL* genes can be divided into two groups: i) those with rather constitutive and stable expression levels during all *Arabidopsis* developmental stages, and ii) those showing high expression levels during specific growth and reproduction processes of *Arabidopsis* development. Similar division can be observed in moss *P. patens* where the expression data clearly show that Pp*SPL* genes from Group 1 are not expressed or very weakly expressed in spores and protonema while in gametophores and sporophyte their expression level is prominent and stable (Fig. 6B). The *PpSPL7* gene showed the highest expression in archegonia and different stages of sporophyte development what may suggest its importance during moss sexual reproduction and sporophyte maturation. The Pp*SPL* genes from Group 2 showed higher expression during premeiotic to meiotic stages of sporophyte development (sporophyte PM-M) with the exception of Pp*SPL3* which additionally showed high expression in rhizoids and chloronema. The other two *Physcomitrium SPL* groups exhibited constitutive expression in all analyzed moss tissues and developmental stages.

As observed in *Arabidopsis* and *Physcomitrium*, also *M. polymorpha* and *A. agrestis SPL* genes belonging to Group 3 and Group 4 exhibited rather constitutive expression profiles in all types of organs and developmental stages analyzed (Fig. 6C and D). In *Marchantia, SPL* members belonging to Group 1 and Group 2 showed rather tissue specific expression with the highest expression observed in reproductive organs development and in young sporophyte. This finding may indicate that the Mp*SPL* genes are involved in the entire process of growth and development in this liverwort with some additional role for Mp*SPL1* and Mp*SPL2* during sexual reproduction as their expression is up-regulated in *Marchantia* sex organs (Fig. 6C). In the case of hornwort *A. agrestis*, the most specific expression pattern was observed for Aa*SPL2* belonging to Group 2 which expression is mostly found in the sporophyte generation while Aa*SPL1* belonging to Group 1 showed equal expression levels during both gametophyte and sporophyte development (Fig. 6D).

The expression data analysis showed that in all analyzed plant species, the *SPL* genes may fall into one of the two categories in the context of expression profile. First one, genes which are highly expressed in nearly all tissues and that is why may function similarly as housekeeping genes for the maintenance of basal cellular function (genes from Group 3 and Group 4). What is more, the genes belonging to this category are not regulated by miRNAs. The second category consists of genes with developmentally specified or enriched expression which are important for the regulation of specific processes during growth and reproduction. Importantly, many genes from this category are under post-transcriptional control guided by miRNA (Fig. 1). In three out of four analyzed plant species, including dicot *A. thaliana*, moss *P. patens* and liverwort *M. polymorpha, SPL* genes which expression profile is strongly correlated with sexual reproduction (genes from Group 1 and group 2) were found. Since there is no data concerning gene expression in the reproductive organs of hornwort *A. agrestis*, based on the observed evolutionary conserved mode of action for some representatives within the *SPL* family, it might be hypothesized that most probably also in *Anthoceros* at least one of the *SPL* family members could be engaged in the regulation of the reproductive pathway.

## Discussion

Bryophytes comprising hornworts, liverworts and mosses are sister group to the vascular plants which can give us information on how their ancestors managed to colonize the terrestrial habitat. In our study, we took an advantage of available genomes of liverwort, *M. polymorpha*, moss *P. patens* and two hornwort species, *A. agrestis* and *A. punctatus* to identify the members of *SPL* gene family and further to investigate their phylogenetic relationships with the angiosperm representative, *A. thaliana. SPL* genes form a major family of plant-specific transcription factors and encode proteins with a highly conserved SBP-box DNA-binding domain. They are crucial players regulating different biological processes in plants, including juvenile to adult phase transition, vegetative to reproductive phase transition, apical dominance, flower development and many more (Chen *et al*., 2010; Xu *et al*., 2016).

No SBP-box related sequences for hornworts were available in the public databases at the time we started our attempt to identify SBP-box genes from this plant lineage. In our study, four *SPL* genes were identified in both hornwort species, *A. agrestis* and *A. punctatus* what is similar to the set of *SPL* genes observed in the liverwort *M. polymorpha* (Tsuzuki et al., 2016; Li et al., 2020). The number of *SPL* gene family members from hornworts and liverworts is in contrast to data published for all other land plants where it is shown that already in moss *P. patens*, there are 13 members of *SPL* family and further *SPL* family expansion has occurred in angiosperm genomes with 23 members found in *Medicago truncatula* (Wang *et al*., 2019), 31 in *Zea mays* (Zhang *et al*., 2016) and even 56 in *Triticum aestivum* (Li et al., 2022b). This shows that the number of *SPL* genes varies considerably between genomes of land plants. The identification of only four *SPL* members in hornworts and liverworts as compared to other land plants underlines that the evolution of all land plant *SPL* genes was a result of several rounds of gene duplication and next speciation events of the paralog genes which originated from only four *SPL* representatives present in the basal lineages of bryophytes. This hypothesis supports the phylogenetic tree of SPL proteins from bryophytes and *Arabidopsis* (Fig. 1) where in each out of four phylogenetic groups, only single representants from *Anthoceros* and *Marchantia* are recognized. Additionally, in the all identified phylogenetic groups, the SPL proteins from bryophytes are grouped on neighboring branches reflecting their close evolutionary relationships.

Gene structure analysis revealed that *SPL* genes from bryophytes and *Arabidopsis* share similar exon-intron organization within the same phylogenetic group with the exception of *Anthoceros SPL* genes from Group 3 and Group 4. Hornworts *SPL* genes from Group 3 and Group 4 possess only one or two very short introns in comparison to the complex structures of *SPL* genes from the liverwort *M. polymorpha*, moss *P. patens* and dicot *A. thaliana* (Fig. 2). Additionally, the average intron length for hornworts *SPLs* is the shortest when compared to *SPL* genes from the remaining land plants used in our study. The specificity of the intron length and number within *SPL* genes in both *Anthoceros* species correlates with the high gene density in their genomes, which is achieved by the presence of many intron-less genes. Additionally, the gene structure of these *SPL* genes reflects a characteristic feature of both hornwort genomes which is the presence of three to four exons per gene on average (‘The Genome of the Model Species Anthoceros agrestis’, 2016; Li *et al*., 2020). On the other hand, the liverwort *SPL* genes show the highest average value for intron length (Fig. 2) what is also in agreement with *Marchantia* average intron length calculated from the genome analysis (Bowman *et al*., 2017). To conclude, evidence based on available genomic data indicates the conservation of exon-intron structures within *SPL* clades with only slight variation in the number of exons and introns mostly observed in hornworts. This conservation is observed even between distantly related species like liverwort *M. polymorpha* and angiosperm *A. thaliana*. However, exceptions to this rule of *SPL* gene structure conservation can be found, like in *Anthoceros*, which can be related to the genome composition and structure.

The four phylogenetic *SPL* families obtained in our study were further analyzed in the context of protein motifs conservation. We observed that the SPL proteins showed a similar pattern of conserved motifs between bryophytes and *Arabidopsis* in Group 2, Group 3 and Group 4, (Fig. 2), with the exception of *Anthoceros* SPL3 proteins. However, in Group 1 the SPL proteins differed explicitly between analyzed plant species. Only the SBP domain was found to be a common motif for all SPL proteins regardless of the land plant lineage. Moreover, very high amino acid conservation was found within the SBP domain, in particular for the zinc-finger like structures and the NLS signal (Fig. 5A-D). As shown in structural studies using *Arabidopsis* SPL proteins, all conserved basic amino acids from Zn-1, Zn-2 and NLS signal form a positively charged surface involved in binding the negatively charged DNA (Yamasaki *et al*., 2004). Although SPL proteins were also described in green algae, *Chlamydomonas* SBP domains showed lower degree of conservation in the amount of basic amino acids, especially within the first zinc-finger like structure (Fig. 5E). In fact, Birkenbihl and co-workers have shown that *C. reinhardtii* CRR1 protein exhibited a significantly lower affinity to the *Arabidopsis*-derived 15 bp *AP1* promoter fragment and to the *Chlamydomonas*-derived copper response element (CuRE) in comparison to *Arabidopsis* AtSPL1, AtSPL3, AtSPL8 and moss PpSPL1 proteins (Birkenbihl *et al*., 2005). Therefore, the lower amount of basic amino acid in the green algae SBP domain of the CRR1 protein when compared to land plants might be responsible for its lower efficiency to interact with DNA. Among the conserved Arg/Lys residues, those in the N-terminal part of the SBP domain (Lys14, Arg/Lys18, Arg19, Lys/Arg21) are suggested to be the candidate residues that determine the sequence specificity by direct recognition of the DNA bases (Yamasaki *et al*., 2004). All these conserved amino acid residues are present in the SBP domains from all three bryophyte species, indicating that those positions were fixed very early during land plants evolution.

Along with the SBP domain, we found additional motifs in the analyzed SPL proteins which especially in Group 3 and Group 4 showed high conservation between evolutionary distant plant species (Fig. 3). The function of these motifs is yet unknown, however, because of their high evolutionary conservation they might be considered as structural units important for proper function of encoded SPL proteins. Based on the analyzed expression profiles, all *SPL* genes from Group 3 and Group 4 exhibited constitutive expression across different organs and developmental stages in the representatives of bryophytes and in *Arabidopsis* and hence, can be important factors for the regulation of basic cellular processes in all land plants. The SBP domain is crucial for specific recognition and binding to *cis*-elements in the promoter of nuclear genes to regulate their expression. However, the additional conservation within the C-terminal part of those proteins may indicate that these conserved motifs are important for the Group 3 and Group 4 SPL proteins to orchestrate the proper expression profile in different tissues and organs throughout the plant life cycle. This could be achieved by interaction of these SPLs with other proteins via conserved C-terminal localized motifs, for example the ankyrin repeats which are known to be involved in protein–protein interactions. Still, the significance of these conserved motifs remains unknown and needs to be further investigated, especially using cross species studies.

The most variable motif composition in the analyzed SPL proteins was observed in Group 1 where additionally, differences in protein lengths were observed (Fig. 3). It seems that during the course of evolution, the *Physcomitrium* Group 1 SPLs diversified independently from other family members in moss lineage leading to their functional specialization. From experimental studies it was shown that Pp*SPL1* and Pp*SPL4* genes regulate proper protonema branching, gametophore development and spore germination (Riese *et al*., 2008). Although PpSPL7 protein possesses the same set of conserved protein motifs as PpSPL1 and PpSPL4, its expression profile may indicate its involvement in archegonia and sporophyte development (Fig. 6B). Thus far, similar motif composition between proteins from the same family is not necessarily related to performing functions in the same developmental processes. Whether the identified *Physcomitrium* Group 1 unique-motifs are crucial to fulfill their specific roles is a matter of future investigations.

The promoter region composition is a key element involved in the regulatory control of gene expression in a tissue specific manner or in response to different stimuli. Many *cis*-elements were found in the promoter regions of *SPL* genes from analyzed bryophytes and *Arabidopsis*, mostly associated with growth and development, light, hormone, and stress responsiveness (Fig. 5). This data indicates that in each of the studied plant species, the *SPL* family is under complex and elaborate control of the transcription, regulated by various environmental and developmental changes. Interestingly, no similar set of *cis*-elements distribution was observed in the promoter regions of *SPLs* within the same phylogenetic group implying that the alteration of *cis*-regulatory elements took place during the land plants *SPL* genes evolution.

In order to further explore the expression landscape of *SPL* genes from the selected plant species, the expression profiles of investigated *SPL* genes were analyzed from different developmental stages and organs of each plant (Fig. 6). The obtained heat maps of expression profiles revealed that both bryophytes and *Arabidopsis SPL* genes from phylogenetic Group 3 and Group 4 exhibit constitutive expression while *SPLs* belonging to Group 1 and Group 2 are expressed in a developmentally specific way or their expression is higher in specific organs/tissues. This differentiated expression pattern correlates with the posttranscriptional expression regulation by miR156 or miR529 family members of all genes from Group 2 and three *Arabidopsis SPLs* from Group 1 (Fig. 1). miR156 is conserved across all land plant lineages while miR529 is mostly present in bryophytes and monocots. Based upon functional and evolutionary analyses, miR529 seems to have been lost in some taxonomic groups, including core eudicots such as *A. thaliana* (Xie *et al*., 2021; Li *et al*., 2022*a*). In *Arabidopsis*, miR156 functions as a master regulator of the juvenile plants development and slows down the vegetative to reproductive phase transition (Chen *et al*., 2010; Sunkar, 2012; Zheng *et al*., 2019). On the other hand, both miRNAs are present in rice and are known to fine-tune the expression of *SPL* genes but at different stages of rice development. First, miR156 represses the *SPL* genes at the vegetative stage to regulate branch outgrowth and next, at the reproductive stage miR529 down-regulates the *SPL* genes to properly define panicle size and architecture (Wang *et al*., 2015; Li *et al*., 2022*a*). Similar expression pattern for both these miRNAs was also observed in moss *P. patens* where miR156 was mainly expressed in protonema while miR529 was mainly expressed in gametophores with developed sporophytes (Xie *et al*., 2021). miR156 and miR529 are also expressed in liverwort *Pellia endiviifolia* although no functional studies are available on their mode of actions in that species (Alaba *et al*., 2015; Pietrykowska *et al*., 2022). In contrast, miR156 is not present in *M. polymorpha* and only miR529c regulates the reproductive transition by repressing Mp*SPL2* gene expression during vegetative growth, thereby preventing the development of reproductive branches and reproductive organs (Tsuzuki *et al*., 2019). Thus, the miR529–*SPL* module seems to control the vegetative-to-reproductive phase change in liverworts development, similarly to miR156/529–*SPL* module in angiosperms. Although we did not find any proof of miR156 and miR529 presence in the genomes of investigated *Anthoceros* species, our analysis revealed that the conserved miR156/529-responsive element in *AaSPL2* and *ApSPL2* genes can be recognized. Thus, it is highly likely that at least one of these miRNAs is present in the investigated *Anthoceros* species, especially since in another *Anthoceros* species, *A. angustus*, miR156 has been identified (Zhang *et al*., 2020).

It is worth noting that in the phylogenetic Group 1 only three *Arabidopsis SPL* genes are under control of evolutionary conserved miR156. Interestingly, *Marchantia* Mp*SPL1* is also regulated by miRNA, however by liverwort specific Mpo-MR-13 (Tsuzuki *et al*., 2016; Streubel *et al*., 2023). Based on transcriptomic studies it was suggested that this Mpo-MR-13– Mp*SPL1* module might be involved in controlling the transition from vegetative to reproductive life cycle. Characteristic expression pattern of Mp*SPL1* has been observed with an explicit expression peak in gametangiophores along with simultaneous down-regulation of Mpo-MR-13 precursors at this developmental stage (Flores-Sandoval *et al*., 2018). However, recent functional studies revealed a role of this Mpo-MR-13–Mp*SPL1* module in the regulation of meristem dormancy with superior control of this module by PIF-mediated phytochrome signaling (Streubel *et al*., 2023). The light signaling modulates the architecture of the thallus branching via Mp*SPL2* action which under shade imitating conditions promotes meristem dormancy by repressing meristem activity (Streubel *et al*., 2023). A similar dependence was proposed in *A. thaliana*, where PIF-mediated repression of several *MIR156* genes expression releases the miR156-taregeted *SPLs* from the posttranscriptional control enabling them to pursue the shade avoidance mechanism (Xie *et al*., 2017). Based on these observations, Streubel and co-workers hypothesized that the miRNA–*SPL* regulation module involved in the meristem dormancy important for the shade avoidance mechanism evolved independently in the liverwort and angiosperm lineages (Streubel *et al*., 2023). On the other hand, the authors only focused on the vegetative thallus development and did not show any data connected with the sexual reproduction (Streubel et al 2023). Therefore, it cannot be excluded that the Mpo-MR-13–Mp*SPL1* module may play a dual role during *Marchantia* life cycle.

## Conclusions

In summary, this study reports for the first time phylogenetic and diversification studies of the *SPL* gene family members from representatives of all three bryophyte classes and the model angiosperm *A. thaliana*. We have identified four *SPL* genes in each of the two hornworts species, *A. agrestis* and *A. punctatus*, what is similar to the four *SPL* genes present in the liverwort *M. polymorpha*. Currently, the *SPL* families from hornworts and liverworts are the smallest known *SPL* families in comparison to other land plants. Therefore, most probably they resemble an archetype of *SPL* genes present in the ancestor of today’s land plants from which all other *SPL* members might have originated. From our analysis we proposed four phylogenetic *SPL* groups common in bryophytes and *Arabidopsis*. Only the SBP domain is a common feature identified for all SPL proteins regardless of the land plant lineage. However, depending on the phylogenetic group, we observed that in three out of four groups the SPL proteins showed a similar pattern of conserved motifs between bryophytes and *Arabidopsis*, while in one group diverse motif composition was found. Moreover, two distinct expression patterns were revealed for the *SPL* family members. We observed that the miRNA-targeted *SPL* genes were expressed in a developmentally specific manner while the non-targeted *SPL* genes exhibited constitutive expression, suggesting their primary role in maintaining basal cellular functions. Our analysis together with the literature data indicate that the miRNA–*SPL* regulatory*--* module appeared very early during land plant evolution. It seems that this miRNA-mediated expression regulation for *SPL* genes from Group 2 is conserved in land plants. Instead, for genes belonging to Group 1, the miRNA mode of regulation evolved independently in the liverwort and angiosperm lineages. In the case of hornworts, in-depth analyses are needed to learn whether the miRNA–*SPL* regulatory*--* modules also function to direct specific developmental traits. Our study emphasizes the importance of research on the biological relevance of *SPL* genes from bryophyte representatives to provide a better understanding of the *SPL* family evolution and function.

## Author Contributions

IS: conceptualization; AA: performed analysis; IS and AA: writing – original draft; ZSK and IS: writing – review & editing.

## References

Alaba S, Piszczalka P, Pietrykowska H, et al. 2015. The liverwort Pellia endiviifolia shares microtranscriptomic traits that are common to green algae and land plants. The New phytologist 206, 352–367.

Bailey TL, Johnson J, Grant CE, Noble WS. 2015. The MEME Suite. Nucleic acids research 43, W39–49.

Birkenbihl RP, Jach G, Saedler H, Huijser P. 2005. Functional dissection of the plant-specific SBP-domain: overlap of the DNA-binding and nuclear localization domains. Journal of molecular biology 352, 585–596.

Bowman JL, Kohchi T, Yamato KT, et al. 2017. Insights into Land Plant Evolution Garnered from the Marchantia polymorpha Genome. Cell 171, 287–304.e15.

Cardon G, Höhmann S, Klein J, Nettesheim K, Saedler H, Huijser P. 1999. Molecular characterisation of the Arabidopsis SBP-box genes. Gene 237, 91–104.

de Castro E, Sigrist CJA, Gattiker A, Bulliard V, Langendijk-Genevaux PS, Gasteiger E, Bairoch A, Hulo N. 2006. ScanProsite: detection of PROSITE signature matches and ProRule-associated functional and structural residues in proteins. Nucleic acids research 34, W362–5.

Chao L-M, Liu Y-Q, Chen D-Y, Xue X-Y, Mao Y-B, Chen X-Y. 2017. Arabidopsis Transcription Factors SPL1 and SPL12 Confer Plant Thermotolerance at Reproductive Stage. Molecular plant 10, 735–748.

Chen X, Zhang Z, Liu D, Zhang K, Li A, Mao L. 2010. SQUAMOSA promoter-binding protein-like transcription factors: star players for plant growth and development. Journal of integrative plant biology 52, 946–951.

Cho SH, Coruh C, Axtell MJ. 2012. miR156 and miR390 regulate tasiRNA accumulation and developmental timing in Physcomitrella patens. The Plant cell 24, 4837–4849.

Crooks GE, Hon G, Chandonia J-M, Brenner SE. 2004. WebLogo: a sequence logo generator. Genome research 14, 1188–1190.

Dai X, Zhao PX. 2011. psRNATarget: a plant small RNA target analysis server. Nucleic Acids Research 39, W155–W159.

Fernandez-Pozo N, Haas FB, Meyberg R, et al. 2020. PEATmoss (Physcomitrella Expression Atlas Tool): a unified gene expression atlas for the model plant Physcomitrella patens. The Plant journal: for cell and molecular biology 102, 165–177.

Flores-Sandoval E, Romani F, Bowman JL. 2018. Co-expression and Transcriptome Analysis of Marchantia polymorpha Transcription Factors Supports Class C ARFs as Independent Actors of an Ancient Auxin Regulatory Module. Frontiers in Plant Science 9.

Gasteiger E, Hoogland C, Gattiker A, Duvaud S ‘everine, Wilkins MR, Appel RD, Bairoch A. 2005. Protein Identification and Analysis Tools on the ExPASy Server. The Proteomics Protocols Handbook, 571–607.

Goodstein DM, Shu S, Howson R, et al. 2012. Phytozome: a comparative platform for green plant genomics. Nucleic acids research 40, D1178–86.

Horton P, Park K-J, Obayashi T, Fujita N, Harada H, Adams-Collier CJ, Nakai K. 2007. WoLF PSORT: protein localization predictor. Nucleic acids research 35, W585–7.

Hu B, Jin J, Guo A-Y, Zhang H, Luo J, Gao G. 2015. GSDS 2.0: an upgraded gene feature visualization server. Bioinformatics 31, 1296–1297.

Hultquist JF, Dorweiler JE. 2008. Feminized tassels of maize mop1 and ts1 mutants exhibit altered levels of miR156 and specific SBP-box genes. Planta 229, 99–113.

Jung J-H, Ju Y, Seo PJ, Lee J-H, Park C-M. 2012. The SOC1-SPL module integrates photoperiod and gibberellic acid signals to control flowering time in Arabidopsis. The Plant journal: for cell and molecular biology 69, 577–588.

Kawamura S, Romani F, Yagura M, et al. 2022. MarpolBase Expression: A Web-Based, Comprehensive Platform for Visualization and Analysis of Transcriptomes in the Liverwort Marchantia polymorpha. Plant & cell physiology 63, 1745–1755.

Klein J, Saedler H, Huijser P. 1996. A new family of DNA binding proteins includes putative transcriptional regulators of theAntirrhinum majus floral meristem identity geneSQUAMOSA. Molecular and General Genetics MGG 250, 7–16.

Kropat J, Tottey S, Birkenbihl RP, Depège N, Huijser P, Merchant S. 2005. A regulator of nutritional copper signaling in Chlamydomonas is an SBP domain protein that recognizes the GTAC core of copper response element. Proceedings of the National Academy of Sciences 102, 18730–18735.

Lang D, Zimmer AD, Rensing SA, Reski R. 2008. Exploring plant biodiversity: the Physcomitrella genome and beyond. Trends in plant science 13, 542–549.

Lescot M. 2002. PlantCARE, a database of plant cis-acting regulatory elements and a portal to tools for in silico analysis of promoter sequences. Nucleic Acids Research 30, 325–327.

Letunic I, Bork P. 2018. 20 years of the SMART protein domain annotation resource. Nucleic acids research 46, D493–D496.

Letunic I, Khedkar S, Bork P. 2021. SMART: recent updates, new developments and status in 2020. Nucleic acids research 49, D458–D460.

Li Y, He Y, Qin T, Guo X, Xu K, Xu C, Yuan W. 2022a. Functional conservation and divergence of miR156 and miR529 during rice development. The crop journal.

Li, Li L, Shi F, et al. 2022b. Conservation and Divergence of SQUAMOSA-PROMOTER BINDING PROTEIN-LIKE (SPL) Gene Family between Wheat and Rice. International Journal of Molecular Sciences 23, 2099.

Li C, Lu S. 2014. Molecular characterization of the SPL gene family in Populus trichocarpa. BMC plant biology 14, 131.

Li F-W, Nishiyama T, Waller M, et al. 2020. Anthoceros genomes illuminate the origin of land plants and the unique biology of hornworts. Nature plants 6, 259–272.

Lin P-C, Lu C-W, Shen B-N, et al. 2016. Identification of miRNAs and Their Targets in the Liverwort Marchantia polymorpha by Integrating RNA-Seq and Degradome Analyses. Plant & cell physiology 57, 339–358.

Liu J, Jung C, Xu J, Wang H, Deng S, Bernad L, Arenas-Huertero C, Chua N-H. 2012. Genome-wide analysis uncovers regulation of long intergenic noncoding RNAs in Arabidopsis. The Plant cell 24, 4333–4345.

Michaely P, Bennett V. 1992. The ANK repeat: a ubiquitous motif involved in macromolecular recognition. Trends in Cell Biology 2, 127–129.

Mistry J, Chuguransky S, Williams L, et al. 2021. Pfam: The protein families database in 2021. Nucleic Acids Research 49, D412–D419.

Miura K, Ikeda M, Matsubara A, Song X-J, Ito M, Asano K, Matsuoka M, Kitano H, Ashikari M. 2010. OsSPL14 promotes panicle branching and higher grain productivity in rice. Nature genetics 42, 545–549.

Moreno P, Fexova S, George N, et al. 2022. Expression Atlas update: gene and protein expression in multiple species. Nucleic acids research 50, D129–D140.

Ortiz-Ramírez C, Hernandez-Coronado M, Thamm A, Catarino B, Wang M, Dolan L, Feijó JA, Becker JD. 2016. A Transcriptome Atlas of Physcomitrella patens Provides Insights into the Evolution and Development of Land Plants. Molecular plant 9, 205–220.

Phytozome v13.

Pietrykowska H, Sierocka I, Zielezinski A, Alisha A, Carrasco-Sanchez JC, Jarmolowski A, Karlowski WM, Szweykowska-Kulinska Z. 2022. Biogenesis, conservation, and function of miRNA in liverworts. Journal of experimental botany 73, 4528–4545.

Posit. 2022. Posit.

Preston JC, Hileman LC. 2013. Functional Evolution in the Plant SQUAMOSA-PROMOTER BINDING PROTEIN-LIKE (SPL) Gene Family. Frontiers in plant science 4, 80.

Quinlan AR, Hall IM. 2010. BEDTools: a flexible suite of utilities for comparing genomic features. Bioinformatics 26, 841–842.

Riese M, Höhmann S, Saedler H, Münster T, Huijser P. 2007. Comparative analysis of the SBP-box gene families in P. patens and seed plants. Gene 401, 28–37.

Riese M, Zobell O, Saedler H, Huijser P. 2008. SBP-domain transcription factors as possible effectors of cryptochrome-mediated blue light signalling in the moss Physcomitrella patens. Planta 227, 505–515.

Shikata M, Koyama T, Mitsuda N, Ohme-Takagi M. 2009. Arabidopsis SBP-box genes SPL10, SPL11 and SPL2 control morphological change in association with shoot maturation in the reproductive phase. Plant & cell physiology 50, 2133–2145.

SIB Swiss Institute of Bioinformatics.

Sigrist CJA, de Castro E, Cerutti L, Cuche BA, Hulo N, Bridge A, Bougueleret L, Xenarios I. 2013. New and continuing developments at PROSITE. Nucleic acids research 41, D344–7.

Sommer F, Kropat J, Malasarn D, Grossoehme NE, Chen X, Giedroc DP, Merchant SS. 2011. The CRR1 Nutritional Copper Sensor in Chlamydomonas Contains Two Distinct Metal-Responsive Domains. The Plant Cell 22, 4098–4113.

Stone JM, Liang X, Nekl ER, Stiers JJ. 2005. Arabidopsis AtSPL14, a plant-specific SBP-domain transcription factor, participates in plant development and sensitivity to fumonisin B1. The Plant journal: for cell and molecular biology 41, 744–754.

Streubel S, Deiber S, Rötzer J, Mosiolek M, Jandrasits K, Dolan L. 2023. Meristem dormancy in Marchantia polymorpha is regulated by a liverwort-specific miRNA and a clade III SPL gene. Current Biology.

Sunkar R. 2012. MicroRNAs in Plant Development and Stress Responses. Springer Science & Business Media.

Swarbreck D, Wilks C, Lamesch P, et al. 2008. The Arabidopsis Information Resource (TAIR): gene structure and function annotation. Nucleic acids research 36, D1009–14.

Szövényi P, Frangedakis E, Ricca M, Quandt D, Wicke S, Langdale JA. 2015. Establishment of Anthoceros agrestis as a model species for studying the biology of hornworts. BMC plant biology 15, 98.

TAIR - Home Page.

Tamura K, Stecher G, Kumar S. 2021. MEGA11: Molecular Evolutionary Genetics Analysis Version 11. Molecular biology and evolution 38, 3022–3027.

The Genome of the Model Species Anthoceros agrestis. 2016. Advances in Botanical Research. Academic Press, 189–211.

Tsuzuki M, Futagami K, Shimamura M, et al. 2019. An Early Arising Role of the MicroRNA156/529-SPL Module in Reproductive Development Revealed by the Liverwort Marchantia polymorpha. Current biology: CB 29, 3307–3314.e5.

Tsuzuki M, Nishihama R, Ishizaki K, Kurihara Y, Matsui M, Bowman JL, Kohchi T, Hamada T, Watanabe Y. 2016. Profiling and Characterization of Small RNAs in the Liverwort, Marchantia polymorpha, Belonging to the First Diverged Land Plants. Plant & cell physiology 57, 359–372.

UZH - Hornworts. Universität Zürich.

Walther D, Brunnemann R, Selbig J. 2007. The regulatory code for transcriptional response diversity and its relation to genome structural properties in A. thaliana. PLoS genetics 3, e11.

Wang H, Lu Z, Xu Y, et al. 2019. Genome-wide characterization of SPL family in Medicago truncatula reveals the novel roles of miR156/SPL module in spiky pod development. BMC genomics 20, 552.

Wang L, Sun S, Jin J, Fu D, Yang X, Weng X, Xu C, Li X, Xiao J, Zhang Q. 2015. Coordinated regulation of vegetative and reproductive branching in rice. Proceedings of the National Academy of Sciences of the United States of America 112, 15504–15509.

Waterhouse AM, Procter JB, Martin DMA, Clamp M, Barton GJ. 2009. Jalview Version 2--a multiple sequence alignment editor and analysis workbench. Bioinformatics 25, 1189–1191.

Website.

Xie Y, Liu Y, Wang H, Ma X, Wang B, Wu G, Wang H. 2017. Phytochrome-interacting factors directly suppress MIR156 expression to enhance shade-avoidance syndrome in Arabidopsis. Nature communications 8, 348.

Xie Q, Wang X, He J, et al. 2021. Distinct evolutionary profiles and functions of microRNA156 and microRNA529 in land plants. International journal of molecular sciences 22, 11100.

Xu M, Hu T, Zhao J, Park M-Y, Earley KW, Wu G, Yang L, Poethig RS. 2016. Developmental Functions of miR156-Regulated SQUAMOSA PROMOTER BINDING PROTEIN-LIKE (SPL) Genes in Arabidopsis thaliana. PLoS genetics 12, e1006263.

Yamaguchi A, Wu M-F, Yang L, Wu G, Poethig RS, Wagner D. 2009. The microRNA-regulated SBP-Box transcription factor SPL3 is a direct upstream activator of LEAFY, FRUITFULL, and APETALA1. Developmental cell 17, 268–278.

Yamasaki K, Kigawa T, Inoue M, et al. 2004. A novel zinc-binding motif revealed by solution structures of DNA-binding domains of Arabidopsis SBP-family transcription factors. Journal of molecular biology 337, 49–63.

Zhang W, Bei LI, Bin YU. 2016. Genome-wide identification, phylogeny and expression analysis of the SBP-box gene family in maize (Zea mays). Journal of Integrative Agriculture 15, 29–41.

Zhang J, Fu X-X, Li R-Q, et al. 2020. The hornwort genome and early land plant evolution. Nature plants 6, 107–118.

Zhang S-D, Ling L-Z, Yi T-S. 2015. Evolution and divergence of SBP-box genes in land plants. BMC genomics 16, 787.

Zhang H, Zhang L, Han J, Qian Z, Zhou B, Xu Y, Wu G. 2019. The nuclear localization signal is required for the function of squamosa promoter binding protein-like gene 9 to promote vegetative phase change in Arabidopsis. Plant molecular biology 100, 571–578.

Zheng C, Ye M, Sang M, Wu R. 2019. A Regulatory Network for miR156-SPL Module in Arabidopsis thaliana. International Journal of Molecular Sciences 20, 6166.

Zhu T, Liu Y, Ma L, Wang X, Zhang D, Han Y, Ding Q, Ma L. 2020. Genome-wide identification, phylogeny and expression analysis of the SPL gene family in wheat. BMC plant biology 20, 420.

